# Cooperatively Designed Aptamer-PROTACs for Spatioselective Degradation of Nucleocytoplasmic Shuttling Protein for Enhanced Combinational Therapy

**DOI:** 10.1101/2022.11.01.514682

**Authors:** Ran Liu, Zheng Liu, Mohan Chen, Hang Xing, Jingjing Zhang

## Abstract

Nucleocytoplasmic shuttling proteins (NSPs) has emerged as a promising class of therapeutic targets for many diseases including cancer. However, most reported NSPs-based therapies largely rely on small molecule inhibitors with limited efficacy and off-target effects. Proteolysis targeting chimera (PROTAC) represents a revolutionary inhibitory modality for targeted protein degradation with many advantages, including substoichiometric catalytic activity, improved selectivity, and high efficacy. However, the majority of reported PROTACs so far are still limited to the degradation of cytoplasmic proteins and lack of tumor-specific targeting. To realize the full potential of the PROTAC technology and broaden its applications for the degradation of NSPs, we herein report a conceptual approach for the design of a new archetype of PROTAC (denoted as PS-ApTCs) by introducing phosphorothioate-modified AS1411 aptamer to a ligand of the CRBN E3 ligase, realizing tumor-targeting and spatioselective degradation with improved efficacy. We have demonstrated that PS-ApTCs is capable of effectively degrading nucleolin in target cell membrane and cytoplasm but not in the nucleus, in a CRBN- and proteasome-dependent manner. In addition, PS-ApTCs exhibits superior antiproliferation, pro-apoptotic, and cell cycle arrest potencies. Importantly, we demonstrate for the first time that combination of PS-ApTCs-mediated nucleolin degradation with aptamer-drug conjugates-based chemotherapy can enable an AND-gated synergistic effect on tumor inhibition. Collectively, our results suggest that PS-ApTCs possess the ability to expand the PROTACs toolbox to even wider range of targets in subcellular localization and accelerate the discovery of new combinational therapeutic approaches.

For Table of Contents Only

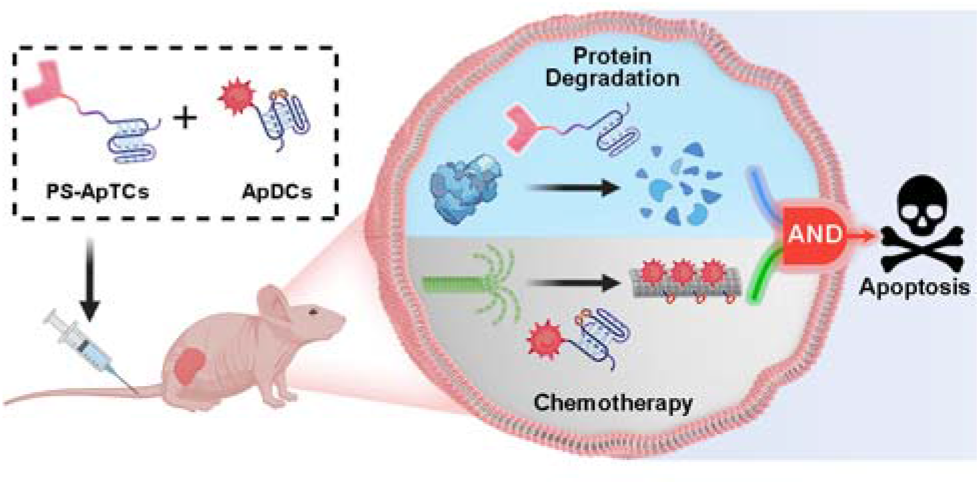

## INTRODUCTION

Nucleocytoplasmic shuttling, one of the major mechanisms to transport a variety of molecules in living cells, plays critical roles in diverse cellular functions.^1^ In particular, the dysregulation of the nucleocytoplasmic shuttling has been found to invoke various diseases including cancer.^2^ For example, the nucleocytoplasmic shuttling proteins (NSPs) that constantly shuttle between the nucleus and cytoplasm play an important role in promoting abnormal cell survival, tumor progression, and drug resistance.^3,4^ Given the importance of NSPs in tumor research, many efforts have been made to develop small molecule-based inhibitors capable of targeting NSPs for potential intervention of nucleocytoplasmic shuttling.^5–7^ Despite significant progress, intrinsic or acquired resistance limits the therapeutic efficacy of small molecule-based inhibitors in patients.^3^ In addition, the “druggable proteome” of NSPs is hampered by the competitive- or occupancy-driven mechanism of traditional inhibitors. Thus, there is still a need for the discovery of new strategies to revert the aberrant nucleocytoplasmic shuttling.

Proteolysis targeting chimeras (PROTACs) are a heterobifunctional class of small molecules that simultaneously recruit a target protein and an E3 ubiquitin ligase complex to trigger target polyubiquitination and subsequent proteasomal degradation.^8–12^ Compared to classical occupancy-based inhibitors, PROTACs exhibit several important advantages, including substoichiometric catalytic activity,^13,14^ improved selectivity,^15,16^ and ability to eliminate the accumulation of drug targets and overcome drug resistance,^17,18^ and potent degrade undruggable targets.^19–23^ As a result, a fast growing number of protein targets have been extensively documented to be degraded by PROTACs.^24–30^ Despite substantial progress, there are still several issues that need to be addressed before its clinical translation is realized.^31^ Firstly, the majority of these systems are limited to small molecule-based PROTACs, which cannot locate more precisely at the target tumor tissue and thus often suffer from the off-target effects. Some progress has been made towards the generation of PROTACs-based conjugates from various functional ligands such as antibodies,^32–35^ folate,^36,37^ or aptamers,^38^ and demonstrated their utility in targeted PROTACs delivery and subsequent tumor-selective degradation of different proteins. However, these methods often involve multiple sophisticated synthesis, including chemical conjugation of the targeting moieties with small molecule-based PROTACs, or embedding cleavable linkers in PROTACs that could be decaged by intracellular stimuli, making them less flexible in warhead design and thus may greatly reduce optimization options of potential drug candidates. Secondly, while most of traditional PROTACs are limited to the degradation of cytoplasmic proteins, the recently developed antibody-based PROTACs,^39^ bispecific aptamer chimeras,^40^ and nanobody-based PROTACs^41^ demonstrated the successful degradation of membrane proteins. However, the PROTAC that permits the spatioselective degradation of protein targets in various cellular compartments has not yet reported due to the lack of a design methodology. Finally, the complexity, diversity, and heterogeneity of tumors has propelled the shift of treatment from monotherapy to polytherapy for enhanced therapeutic outcome. Such a concept also has great significance for the future development and clinical translation of PROTACs but has rarely been explored.

With these considerations in mind, herein we present a conceptual approach for the design of a new archetype of PROTACs (denoted as PS-ApTCs) by introducing phosphorothioate-modified aptamers to an E3 ligand (Scheme 1A). Different from previous aptamer-PROTAC conjugates, PS-aptamer serves as a versatile scaffold for both cell-specific PROTACs delivery and efficient recruitment of target protein. Using nucleolin (NCL) as a model NSP, we demonstrated that PS-ApTCs can recruit E3 ligase CRBN to nucleolin in human cervical cancer cells and potently induce the degradation of nucleolin *in vitro* and *in vivo* (Scheme 1B). Moreover, the anti-tumor superiority of PS-ApTCs were further highlighted by integrating PS-ApTCs with aptamer-drug conjugates-based chemotherapy (Scheme 1C). In addition, our PS-ApTCs provides good versatility over previous aptamer-based PROTACs and novel modalities to overcome current limitations of PROTAC-based monotherapy (Figure S1), as well as new insights into membrane protein degradation (Figure S2). Since *in vitro* selection can obtain aptamers selective for many NSP targets, such a conceptual PROTACs design might also possess the ability to expand the PROTACs toolbox to even wider range of targets in subcellular localization and accelerate the discovery of new combinational therapeutic approaches.

**Scheme 1.**
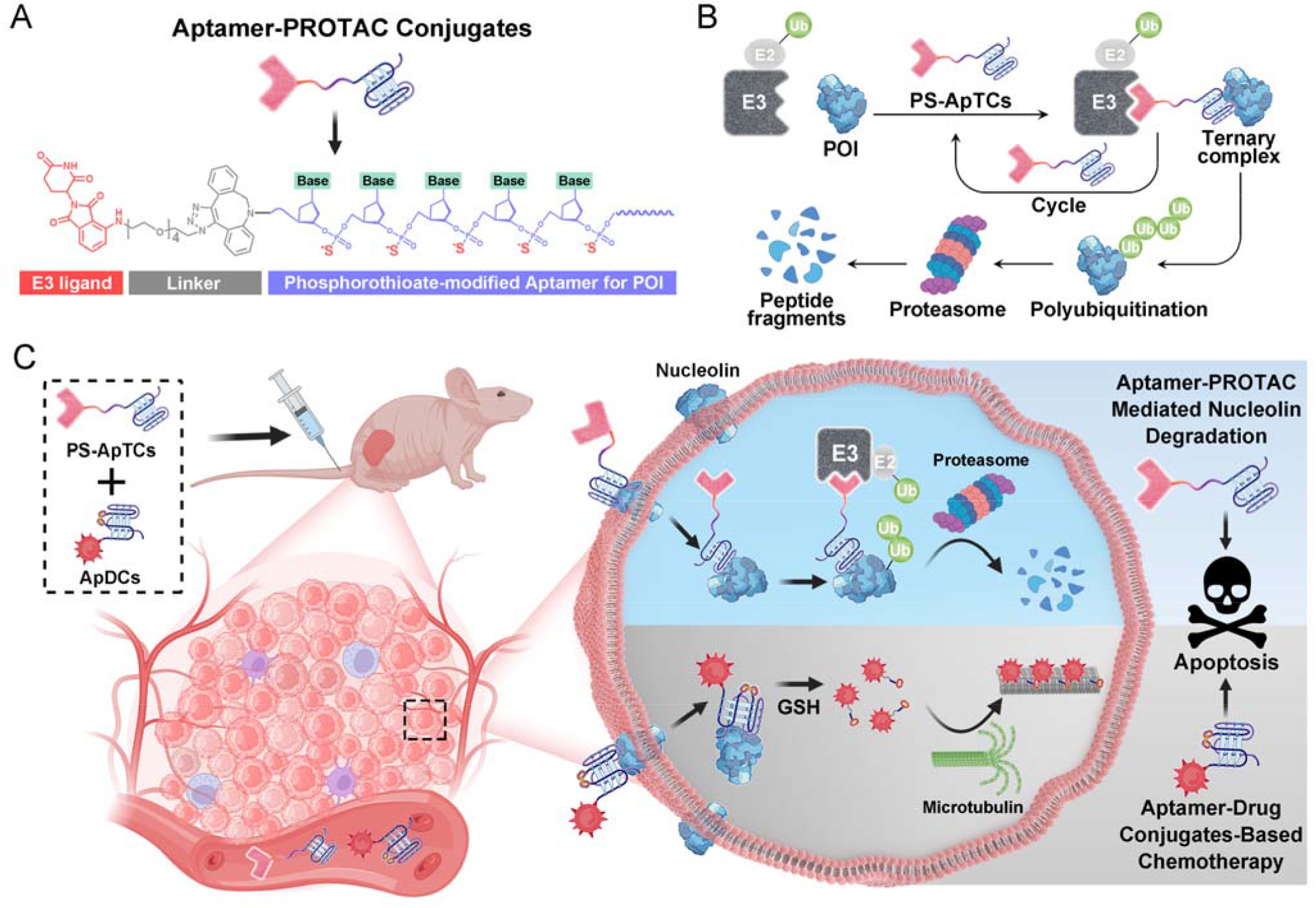
PS-ApTCs-based PROTAC Strategy for Enhanced Combinational Therapy^*a*^. ^***a***^(A) Rational design of PS-ApTCs, which contains an E3 ligand, a linker and a phosphorothioate modified aptamer for targeting proteins of interest (POI). (B) PS-ApTCs simultaneously recruit the E3 ubiquitin ligase to ubiquitinate the POI, which is subsequently degradated by the 26S proteasome. (C) Combination of PS-ApTCs-mediated nucleolin degradation and aptamer-drug conjugates-based chemotherapy for enhanced tumor therapy.

## RESULTS AND DISCUSSION

### Rational Design and Characterization of Phosphorothioate-Modified Aptamer-PROTACs (PS-ApTCs)

A typical PS-ApTCs is composed of three components: a phosphorothioate-modified aptamer warhead to recruit the protein of interest (POI), a ligand to recruit the CRBN E3 ubiquitin ligase, and a linker between these two fragments (Scheme 1A). The critical difference between traditional small-molecule PROTAC and PS-ApTCs is the aptamer targeting warhead enables the PS-ApTCs to specifically recognize target cancer cells, while PS modification enhances stability of PS-ApTCs from nuclease-mediated degradation. These features are generally considered to be a requirement for improving the therapeutic index.^42,43^ To evaluate this experimentally, we first chose the NCL as the POI, and AS1411 aptamer as the NCL-binding domain (Scheme S1A). The former is a shuttling protein overexpressed in cytoplasm and on the cell surface of various cancer cells, while the latter is a well-defined guanine-rich aptamer and has been extensively validated as a highly specific NCL binder *in vitro* and *in vivo*.^42,44–46^ Meanwhile, to ensure that the PS-ApTCs design is generally applicable, pomalidomide, one of the most extensively used CRBN recruiting ligands,^47^ was installed in PS-ApTCs (Scheme S1A). Thus, PS-ApTCs are capable of specifically recruiting both the NCL and the E3 ubiquitin ligase to form a ternary complex. The propinquity allows E3 ubiquitin ligase to induce the polyubiquitination of the NCL and lead to the protein degradation by proteasome (Scheme 1B). In addition, once ubiquitination of the target is completed, the ubiquitinated NCL dissociate from the ternary complex so that the PS-ApTCs can be recycled to reinitiate a new round of recruitment and ubiquitination.

The PS-ApTCs was synthesized through a copper-free click reaction to conjugate commercially available azide-modified pomalidomide (pom-PEG4-azide) to DBCO-modified PS-AS1411 (Figure S3A). The molecular weight from MALDI-TOF MS is similar to the theoretical value (10326.9) (Figure S3B), indicating the successful synthesis of PS-ApTCs. Furthermore, ApTCs, a negative control, was designed and synthesized by replacing the DBCO-modified PS-AS1411 with a DBCO-modified AS1411 (Scheme S1B). Then, we evaluated the nuclease stability of these PROTACs by treating them in 10% fetal bovine serum (FBS) and analyzed using polyacrylamide gel electrophoresis (PAGE). As shown in Figure 1A, the bands corresponding to PS-ApTCs could be observed even after 24 hours of incubation with 10% FBS, whereas the bands corresponding to the ApTCs completely disappeared after 12h. In consistency, the serum half-lives of PS-ApTCs and ApTCs was calculated to be 32 h and 5 h, respectively. Surprisingly, PS-ApTCs also exhibited higher binding affinity to NCL than ApTCs, as confirmed by Bio-Layer Interferometry (BLI) (Figure 1B). Indeed, binding kinetics measurements further confirmed that PS-ApTCs displayed approximately 15-fold greater binding affinity (K_D_=8.7 nM) than that of wild-type AS1411 aptamer (Figure 1C).^48^ These results indicated that PS-modification plays a vital role in improving the serum stability and binding affinity of the PROTACs.

**Figure 1.**
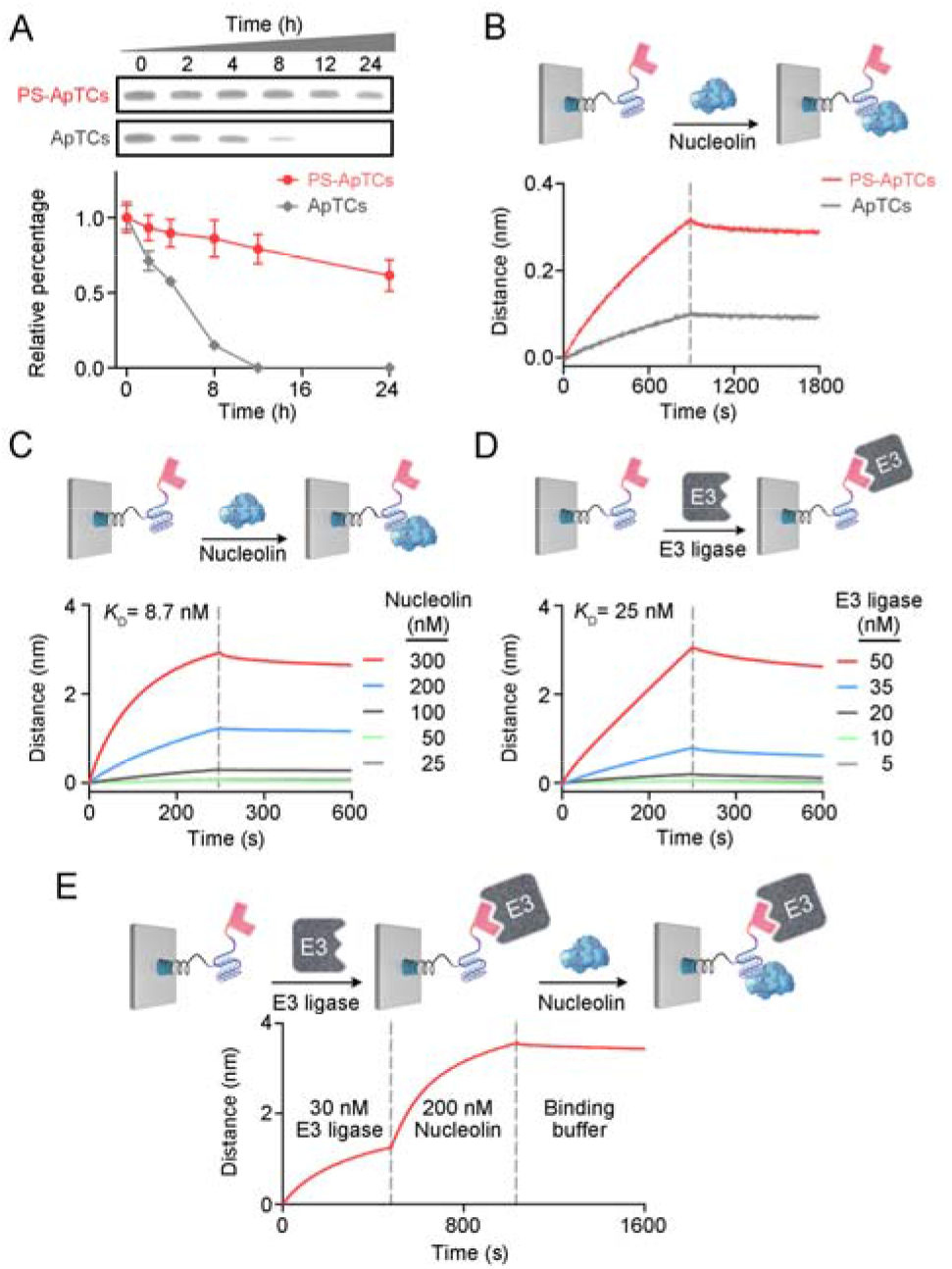
Characterization of PS-ApTCs. (A) Native polyacrylamide gel analysis of the stability of PS-ApTCs and ApTCs in 10% FBS for different times. Quantitative degradation curves showing PS-ApTCs has a significantly prolonged serum half-life. Error bars represent the standard deviations of three independent measurements. (B) BLI experiment showing PS-ApTCs displays higher binding affinity to NCL than ApTCs. Quantitative BLI analysis showing that PS-ApTCs binds to NCL and CRBN with a K_D_ of 8.7 nM (C) and 25 nM (D), respectively. (E) Two-step BLI experiment showing that PS-ApTCs can simultaneously bind to both CRBN and NCL. Biotin-modified PS-ApTCs and ApTCs were captured by streptavidin BLI probes.

Having confirmed the NCL-specific binding, we next examined the binding affinity of PS-ApTCs to CRBN by immobilizing biotinylated PS-ApTCs onto the BLI probe. The BLI binding signals increased in a dose-dependent manner with a K_D_ of 25 nM (Figure 1D), indicating that PS-ApTCs retained high affinity binding to CRBN.^49^ Furthermore, to investigate the feasibility of PS-ApTCs-mediated ternary complex formation, we conducted a similar BLI test by successive incubation in CRBN and NCL solutions. As expected, two successive increases in BLI signal was observed, confirming that PS-ApTCs can recruit both CRBN and NCL concurrently (Figure 1E). Given that the linker length is essential for the formation of sufficiently stable ternary complexes,^50^ we further synthesized a new NCL-recruiting PROTAC degrader (PS-ApTCs-PEG1) using another commercially available azide-modified pomalidomide with a shorter PEG linker length (pom-PEG1-azide) following the same protocol (Figure S4). As shown in Figure S5, compared with PS-ApTCs, PS-ApTCs-PEG1 showed a similar binding affinity to the CRBN, but slightly weaker affinity for the NCL in the two-step BLI experiment. This is likely because a CRBN occupying the binding site of PS-ApTCs-PEG1 may compete with subsequent NCL binding by higher steric hindrance in the presence of a shorter PEG chain length. As a result, the PEG linker with flexible long chain as the bridge between the aptamer and pom-based ligand could not only help to recruit both CRBN and NCL concurrently, but also greatly enhance the freedom of aptamer to improve targeted cell recognition for internalization.

### PS-ApTCs Preferentially Degrades Membrane and Cytoplasmic Nucleolin in Targeted Cancer Cells

Encouraged by the excellent *in vitro* performance of PS-ApTCs, we next explored the cell-specific targeting of PS-ApTCs by incubating FAM-labeled PS-ApTCs with membrane NCL-positive HeLa cells and NCL-negative L-02 cells at 4°C for 60 min, respectively. Confocal fluorescence microscopy images demonstrated that bright green fluorescence was distributed evenly along the HeLa cell membrane, while negligible green fluorescence was observed in L-02 cell membrane following an identical protocol (Figure S6A). These observations were further confirmed by flow cytometric analysis (Figure S6B), which are consistent with those of cells treated with FAM-labeled AS1411 (Figure S6C and S6D). Together, these results demonstrate that the high NCL-targeting function of the AS1411 was maintained for the PS-ApTCs. Subsequently, the time-dependent internalization and intracellular localization of PS-ApTCs was evaluated by confocal fluorescence microscopy. To determine the cellular distribution, CytoTrace red and Hoechst 33342 were used as marker for cytoplasm and nuclei, respectively. As shown in Figure S6E, FAM-labeled PS-ApTCs (green fluorescence) rapidly accumulated in the cell membrane after 10 min incubation, and then started to enter and disperse throughout the cytoplasm at 60 min. Further increasing the exposure time to 240 min resulted in a bright green fluorescence in the cytoplasm as confirmed by colocalization with the red fluorescence from the cytoplasm stain. The Mander’s overlap colocalization coefficient (MOC) showed a positive correlation with the exposure time in the range of 10–240 min. It is noteworthy that a small population of PS-ApTCs still enters the nucleus at 240 min as confirmed by colocalization with the blue fluorescence from the nuclear stain Hoechst 33342 (MOC=0.43). These findings indicated that PS-ApTCs internalized in HeLa cells through NCL-mediated nucleocytoplasmic shuttling process.^51^

We next asked whether PS-ApTCs can be used to degrade NCL in cells. Given that NCL was extensively located in the cell membrane, cytoplasm, and nucleus in cancer cells,^2^ we first examined membrane NCL degradation by harvesting HeLa cells with 200 nM PS-ApTCs at different time points ranging from 0 to 24 h. The membrane NCL was then stained with FAM-AS1411 at 4□. Figure 2A showed that membrane NCL levels decreased in a time-dependent fashion during PS-ApTCs incubation. Quantitative analysis using flow cytometry further confirmed the time-dependent decrease of membrane NCL levels by PS-ApTCs (Figure 2B). For comparision, two negative controls, including a PROTAC synthesized using a randomized DNA sequence (R-PROTAC) and PS-AS1411 (Scheme S1B), were also designed and tested with an identical protocol. Successful synthesis was confirmed through MALDI-TOF MS (Figure S7). Both fluorescence microscopy and flow cytometry showed minimal change in membrane NCL levels when HeLa cells were subjected to either R-PROTACs (Figure S8) or PS-AS1411 (Figure S9). These results indicated that both components of PS-AS1411 and pomalidomide are necessary for the overall function of the PS-ApTCs degrader.

**Figure 2.**
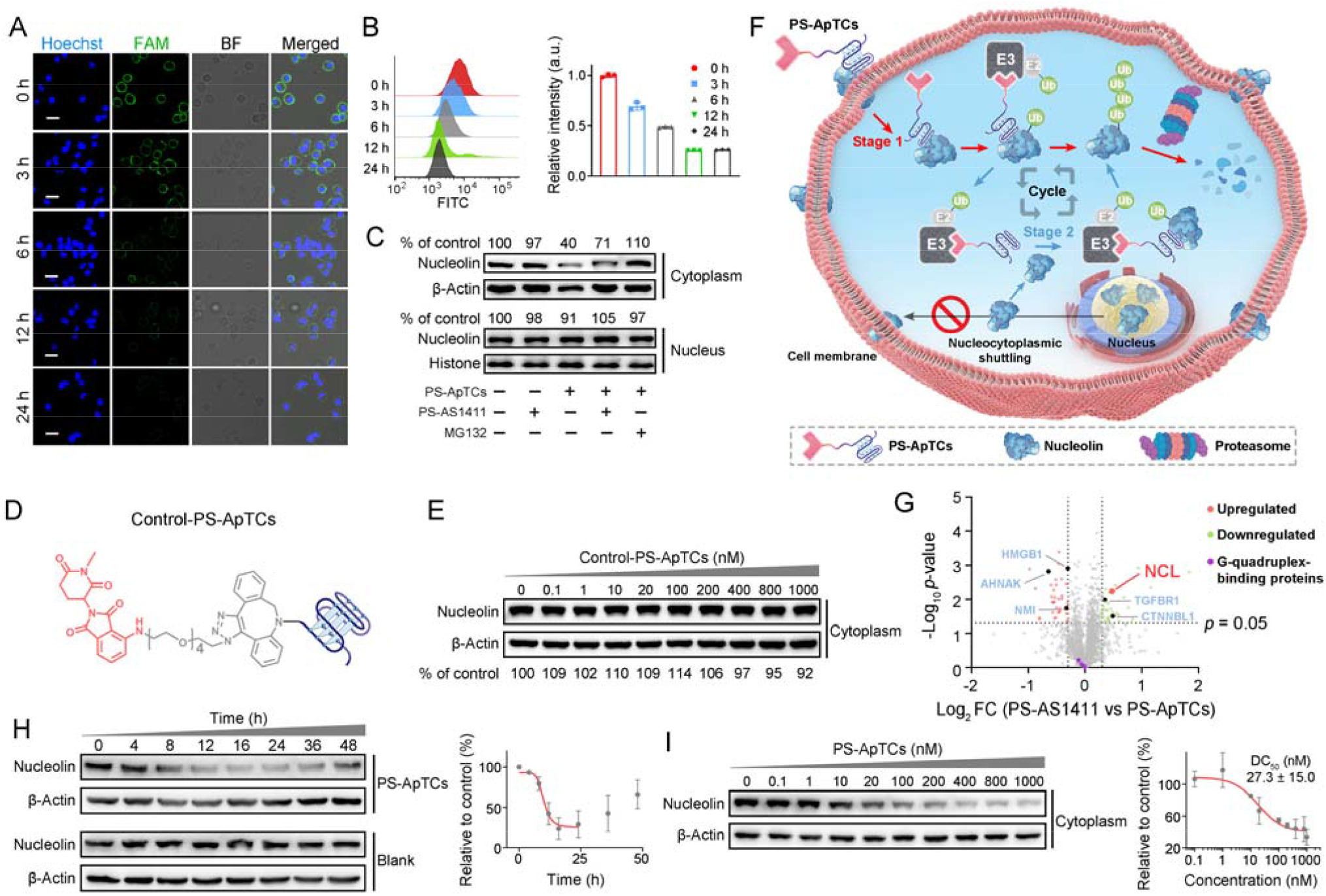
PS-ApTCs efficiently degraded membrane and cytoplasmic NCL in targeted cancer cells. Confocal microscopy images (A) and flow cytometry analysis (B) showing a time-dependent decrease of membrane NCL levels in HeLa cells post-incubation with 200 nM PS-ApTCs. Membrane NCL was labeled with FAM-AS1411 (green) at 4LJ for 60 mins. The scale bar is 20 μm. (C) Western blot analysis showing PS-ApTCs preferentially degraded NCL in cytoplasm but not in the nucleus. (D) Chemical structures of Control-PS-ApTCs. (E) Western blot analysis of cytoplasmic NCL levels in HeLa cells treated with increasing concentrations of Control-PS-ApTCs for 24 h. (F) Schematic representation of PS-ApTCs for targeted degradation of NCL in cell membrane and cytoplasm. (G) Quantitative MS-based proteomics comparing protein abundance profiles of HeLa cells treated with PS-AS1411 or PS-ApTCs (200 nM) for 24 h. (H) Quantitative Western blot analysis showing a time-dependent degradation of cytoplasmic NCL in HeLa cells post-incubation with 200 nM PS-ApTCs. (I) Dose-dependent experiment showing PS-ApTCs efficiently degraded cytoplasmic NCL after 24 h treatment, with a DC_50_ of 27.3±15.0 nM. β-Actin served as a control protein. Error bars represent the standard deviations of three independent measurements.

While we observed potent membrane NCL degradation by PS-ApTCs, it was still not clear how the PS-ApTCs contributes to the degradation process, and more importantly, whether it would degrade NCL in other subcellular components. To answer these questions, we further evaluated the NCL levels in cytoplasm and nucleus of HeLa cells post-incubation with 200 nM PS-ApTCs for 24h. Results by Western blot analysis indicated that cytoplasmic NCL level in PS-ApTCs treated cells was significantly reduced to about 40% compared to untreated control, while only a minimal change of nuclear NCL level was observed (Figure 2C). This result may be attributed to the less effective proteasome degradation of NCL in the nucleus where additional steps or factors are required.^22^ In addition, we found that PS-AS1411 alone was incapable of degrading NCL in either cytoplasm or nucleus, while cotreatment of PS-ApTCs with PS-AS1411 partially inhibited the degradation of cytoplasmic NCL. This observation can be attributed to the competitive internalization of PS-ApTCs and PS-AS1411 through NCL-mediated process.^52^ Notably, the degradation ability of PS-ApTCs was not significantly affected with a shorter PEG chain (Figure S10). Moreover, the degradation of cytoplasmic NCL by PS-ApTCs could be completely blocked by the proteasome inhibitor MG132 (Figure 2C). In addition, we also prepared Control-PS-ApTCs containing a methylated glutarimide ring (Figure S11-S18), which greatly reduces the affinity for CRBN5 and retains the identical NCL binding moiety and linker as PS-ApTCs (Figure 2D). Western blot analysis confirmed that Control-PS-ApTCs had minimal effects on NCL degradation compared to PS-ApTCs (Figure S19A). In addition, we found that Control-PS-ApTCs did not reduce the cytoplasmic NCL level in HeLa cells at concentrations up to 1000 nM for 24 h (Figure 2E). Finally, MLN4924, a commercially-available neddylation inhibitor that disrupts CRBN E3 ligase function,^53–55^ could effectively restore the cytoplasmic NCL level in the PS-ApTCs-treated HeLa cells (Figure S19). Overall, these findings illustrate a possible mechanism of spatioselective degradation of nucleocytoplasmic shuttling NCL (Figure 2F): PS-ApTCs selectively binds to the membrane NCL on tumor cells and the resulting PS-ApTCs-NCL complex is internalized and recruit the CRBN E3 ligase to ubiquitinate the NCL, which is subsequently degraded by the proteasome (Stage 1). After that, PS-ApTCs is recycled to recruit the cytoplasmic NCL and trigger a new round of degradation (Stage 2). These two stages cooperatively facilitate the spatioselective proteolysis of NCL through the disruption of nucleocytoplasmic NCL shuttling.

To assess whether the expression levels of other cytoplasmic proteins are affected by PS-ApTCs, we next performed quantitative dimethyl-based proteomic studies. Briefly, the proteomic samples were obtained by treating HeLa cells with PS-ApTCs, or PS-AS1411 at 200 nM for 24 h, followed by extraction of cytoplasmic proteins. Over 2000 proteins were quantified, and 43 out of 2064 (2.08%) proteins were downregulated and 36 out of 2064 (1.74%) proteins were upregulated by the PS-ApTCs-treated group (Figure 2G), when compared with the control group (PS-AS1411). Moreover, NCL was one of the most significantly downregulated protein, consistent with the immunoblotting results (Figure 2C). Notably, the 10 proteins with a significantly reduced expression level in PS-ApTCs-treated samples, were closely involved in NCL signaling network/pathway (Figure S20), including the cell proliferation modulator of TGFBR1 and CTNNBL1.^56,57^ Meanwhile, the 23 upregulated proteins associated with cell cycle arrest, antiproliferation or apoptosis, including HMGB1, AHNAK, and NMI,^58–61^ were found to have functional relationship with tumor suppressor p53 (Figure S20), indicating a potential p53-dependent antitumor mechanism. Moreover, the PS-ApTCs treatment did not lead to a significant change in protein levels of other G-quadruplex-binding proteins such as CIRBP, RHAU, and SLIRP (Figure 2G).^62–64^ These results suggest that PS-ApTCs is relatively selective for NCL degradation.

To evaluate the degradation efficiency of PS-ApTCs, we further treated HeLa cells with 200 nM PS-ApTCs for different times. Figure 2H showed the time-dependent degradation profiles of cytoplasmic NCL, and we found that the degradation could begin as early as 4 h and plateau at 24 h, giving a lowest cytoplasmic NCL level of 29.2±16.6 % of the initial value. Notably, a trend toward recovery of cytoplasmic NCL was also observed after 24 h, indicating that PS-ApTCs-induced sustained degradation is also dependent on continued exposure of PS-ApTCs. Furthermore, we also found that PS-ApTCs could effectively degraded cytoplasmic NCL in a dose-dependent manner, with a DC_50_ of 27.3±15.0 nM after only 24 h of treatment (Figure 2I), which is comparable to that of previous PROTACs. However, no significant degradation of cytoplasmic NCL was observed in L-02 cells treated with PS-ApTCs for up to 48h (Figure S21), indicating the targeted degradation of NCL by PS-ApTCs in membrane NCL-positive cells. These findings together confirmed that PS-ApTCs effectively and preferentially degraded membrane and cytoplasmic NCL in a CRBN- and proteasome-dependent manner.

### PS-ApTCs Shows Enhanced Antiproliferation, Pro-Apoptotic and Cycle Arrest Potencies

Given that NCL is a promising target for anti-cancer therapy, we further examined the anti-proliferative activity of PS-ApTCs using the CCK-8 assay. As shown in Figure 3A, 200 nM PS-ApTCs displayed a time-dependent response in inhibiting HeLa cell viability during 48□h post-incubation, while the same concentration of R-PROTAC, PS-AS1411, and Control-PS-ApTCs exhibited no antiproliferative activity. Similarly, compared with above three control groups, PS-ApTCs showed an enhanced inhibition of cell proliferation in a dose-dependent manner, with an IC_50_ value of 33.7 nM (Figure 3B). These results suggest that the antiproliferation activity of PS-ApTCs in HeLa cells is likely due to its CRBN-dependent NCL protein degradation activity. In addition, the effect of PS-ApTCs on HeLa cell apoptosis was investigated by flow cytometry analysis with the dual fluorescence of Annexin V-FITC/PI (Figure 3C). For all the PROTACs we tested, PS-ApTCs promoted the largest population of HeLa cells to undergo early- and late-stage apoptosis (Figure 3D), and the result was in accordance with the antiproliferation test. Next, we sought to ask whether NCL degradation affects cell cycle. Figure 3E showed the flow cytometry analysis of cell cycle distribution in HeLa cells treated with different PROTACs as indicated. We found that only PS-ApTCs (200 nM) resulted in cell cycle arrest by increasing the accumulation of HeLa cells in G2/M phase while reducing G0/G1 and S phase cells (Figure 3F). For comparision, when NCL-negative L-02 cells were tested, the PS-ApTCs treatment failed to decrease cell viability (Figure S22A), and all three PROTACs as indicated displayed no obvious inhibition effect even at a high concentration of 400 nM (Figure S22B). In addition, there was no significant effect on apoptosis activation (Figure S23) or cell cycle distribution (Figure S24) for all groups tested, indicating a tumor cell-specific effect by the PS-ApTCs degrader. Collectively, these findings provided significant support for the enhanced antiproliferation, pro-apoptotic and cycle arrest potencies of the rationally designed PS-ApTCs.

**Figure 3.**
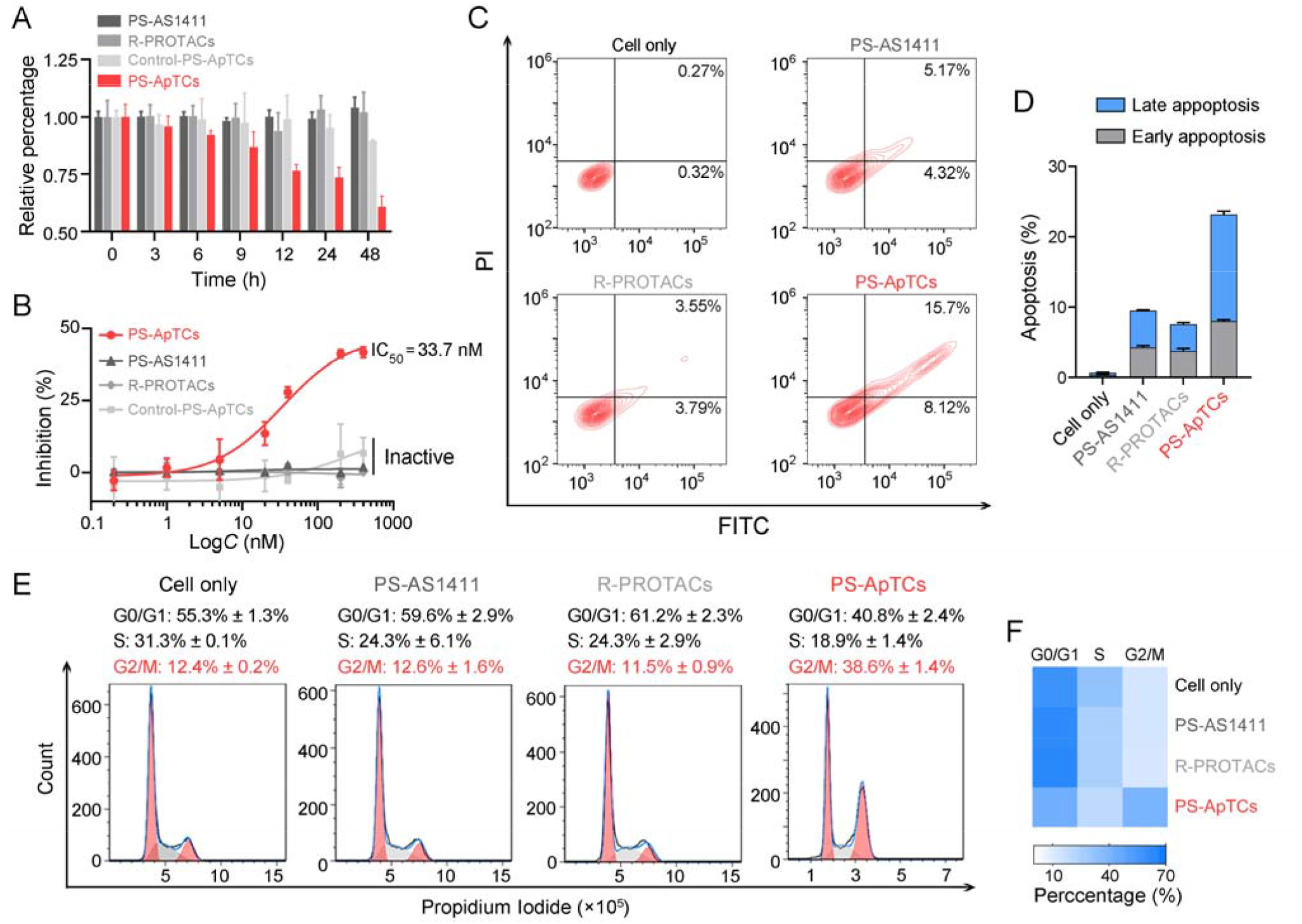
PS-ApTCs showed enhanced antiproliferation, pro-apoptotic and cycle arrest potencies. (A) Cell viability of HeLa cells treated with 200□nM of different PROTACs as indicated for 0 h, 3 h, 6 h, 12 h, 24 h or 48 h in cell culture medium at 37°C. (B) Quantitative dose-dependent experiment showing PS-ApTCs has excellent antiproliferation activity with an IC_50_ of 33.7 nM. HeLa cells were treated with different concentrations of PROTACs as indicated for 48 h. (C) Flow cytometry analysis of HeLa cells post-incubation with 200 nM of different PROTACs for 48 h, followed by staining with Annexin V-FITC, and propidium iodide. (D) Quantitative analysis of apoptosis showing PS-ApTCs induced the highest level of early- and late-stage apoptosis. Error bars represent the standard deviations of three independent measurements. (E) Cell cycle analysis of HeLa cells treated with 200□nM of different PROTACs, as determined by flow cytometry using propidium iodide staining. (F) Representative heatmaps of cell cycle distribution from the assays presented in (E) showing only PS-ApTCs resulted in cell cycle arrest at G2/M phase.

### Design and Characterization of GSH-Responsive AS1411 Aptamer-Drug Conjugates (ApDCs)

Polytherapy, which combines different therapeutic agents to simultaneously disrupt multiple antitumor mechanisms, has been expected as a promising strategy for enhancing therapeutic efficiency in cancer treatment.^65^ The exceptional *in vitro* behaviors of the PS-ApTCs motivated us to further investigate whether the anticancer activity triggered by the PS-ApTCs could be combined with other therapeutic approaches to realize systemic cancer management. As a proof-of-principle experiment, aptamer-drug conjugates (ApDCs), an innovative targeted drug delivery strategy for enhancing the efficacy of chemotherapy,^66^ was chosen for an initial investigation. As illustrated in Figure 4A, the typical ApDCs is composed of three modules: 1) a DBCO-modified AS1411 with incorporation of a reduction-cleavable disulfide bond (−S−S−), 2) an azide-modified paclitaxel (PTX) that acts as an antitumor drug, and 3) a linker containing a hydrophilic PEG segment (Mw=1000). Considering that GSH levels are relatively low in the extracellular matrix (∼2 μM in plasma) but high in the cytoplasm (2−10 mM),^67^ we reasoned that PS-ApTCs could remain in the −S−S− state before it enters the cells, enabling efficient NCL receptor-mediated uptake into the cytoplasm. Moreover, it can be subsequently cleaved by intracellular GSH to release the active PTX for chemotherapy, and more importantly, disrupt G-quadruplex structure of AS1411 to minimize the competition with PS-ApTCs for intracellular NCL occupancy when polytherapy was applied. The proposed ApDCs was synthesized *via* the copper-free click reaction (Figure S25A), and confirmed by mass spectrometry (Figure S25B). Prior to undertaking intracellular evaluation of ApDCs, it was important to first characterize its GSH-responsive property. To this aim, we conducted a PAGE-based cleavage assay in the presence of GSH (2 mM) to mimic the reducing environment. As shown in Figure S26A, GSH-treated DBCO-AS1411-intSH (25−50 bp) results in two DNA fragments (< 25 bp) in perfect agreement with the expected cleavage products upon disulfide bond cleavage in the GSH-responsive aptamer, confirming GSH-responsive releasing feature. In addition, the release of PTX from GSH-treated ApDCs was verified by high-performance liquid chromatography (HPLC). As shown in Figure S26B, ApDCs, DNA fragment-1 and fragment-2 exhibited monodisperse peak at elution time of 18.1, 14.3 and 11.2 min, respectively. After incubation of ApDCs with 2 mM GSH for 1 h, the original peak of ApDCs decreased, while two new peaks with retention times at 14.3 min and 11.1 min appeared, which could be assigned to DNA fragment-1 and PTX-fragment 2, respectively. These results indicate that the ApDCs retains GSH-responsive property of the DBCO-AS1411, liberating the PTX for the subsequent chemotherapy.

**Figure 4.**
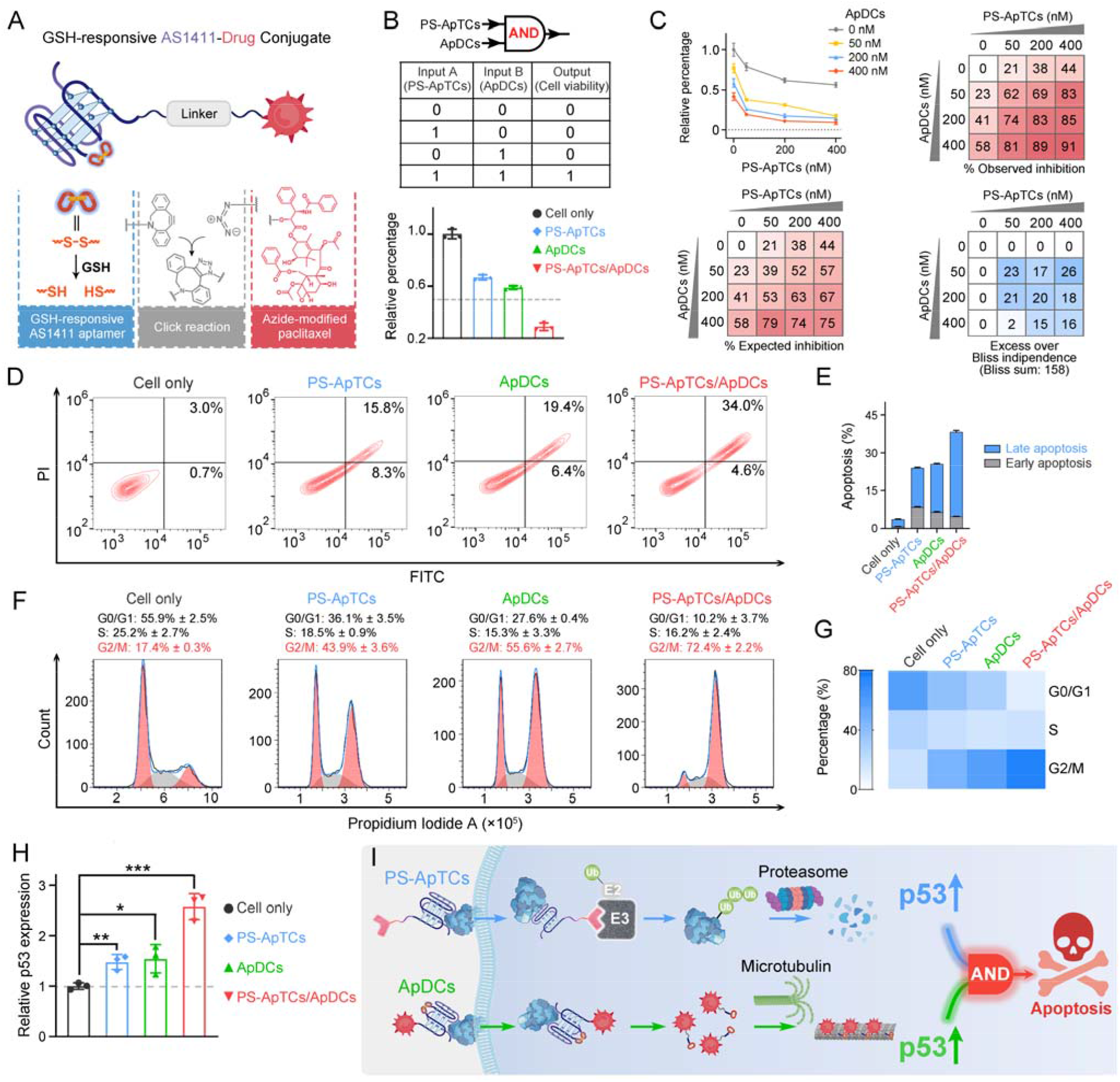
Cooperativity of PS-ApTCs and ApDCs resulted in AND-gated therapeutic effect by upregulating p53. (A) Design of GSH-responsive aptamer-drug conjugates (ApDCs), which incorporates an AS1411 aptamer containing a cleavable disulfide bridge into an azide-modified paclitaxel *via* a copper-free click reaction. (B) Cell viability analysis showing a typical Boolean AND logic function operating with 200 nM of PS-ApTCs and ApDCs for 24 h. A 50% antiproliferation activity was defined to separate the ON and OFF logic states in the AND truth table. (C) Drug synergy test of PS-ApTCs and ApDCs using Bliss independence model. HeLa cells were treated with PS-ApTCs and ApDCs alone and in combination at the indicated concentrations for 24 h. Drug synergy represented by excess over Bliss scores were calculated based on the Bliss Independence model at each combination of drug doses. (D) Flow cytometry analysis of HeLa cells post-incubation with 200 nM of different compounds as indicated for 48□h, followed by staining with Annexin V-FITC, and propidium iodide. (E) Quantitative analysis of apoptosis showing PS-ApTCs and ApDCs displayed apparent positive cooperativity for inducing late-stage apoptosis. (F) Cell cycle analysis of HeLa cells treated with 200□nM of different compounds, as determined by flow cytometry using propidium iodide staining. (G) Representative heatmaps of cell cycle distribution from the assays presented in (F). (H) Quantitative reverse transcription-PCR study showing PS-ApTCs and ApDCs displayed apparent positive cooperativity for upregulating the p53 mRNA expression level in HeLa cells. Error bars represent the standard deviations of three independent measurements. (I) Schematic illustration of cooperatively designed AND-gated PS-ApTCs and ApDCs for enhanced therapeutic purpose.

Having demonstrated the feasibility of GSH-responsive ApDCs, we sought to determine whether ApDCs can affect the ability of PS-ApTCs to induce NCL degradation in the cytoplasm. To investigate this, we performed quantitative analysis of cytoplasmic NCL levels in HeLa cells treated with equivalent concentrations (200 nM) of ApDCs, PS-ApTCs, or PS-ApTCs/ApDCs for 24 h by Western blotting (Figure S27). Compared with untreated cells, ApDCs resulted in no appreciable degradation of cytoplasmic NCL. Meanwhile, we found that PS-ApTCs/ApDCs codosed could effectively degrade the cytoplasmic NCL (>50%), as efficiently as PS-ApTCs. These results indicated that ApDCs had a negligible impact on the degradation activity of PS-ApTCs, enabling potential biorthogonal applications of these two distinct agents for enhanced therapy.

### PS-ApTCs and ApDCs Possess AND-Gated Therapeutic Effect by Upregulating p53

Given that PS-ApTCs and ApDCs undergo the same NCL-mediated internalization, we next determined if these two therapeutic agents could be used to construct logic circuits for enhanced therapy. For this purpose, HeLa cells were treated with PS-ApTCs, ApDCs or their mixture at an equal molar ratio for 24 h, and cell viability was examined using the CCK-8 assay. As depicted in Figure 4B, in a typical assay using 200 nM of PS-ApTCs or ApDCs as the inputs, a 50% cell viability was defined to separate the OFF and ON logic states of the output antiproliferative effect. The antiproliferative effect was distinctly high (output=1) only when the input was in a (1/1) state, while the output was 0 when inputs were (0/0), (0/1), and (1/0) (Figure 4B), which indicates the successful construction of the Boolean AND gate that is “ON” only when both inputs are “ON”. These results suggest that PS-ApTCs and ApDCs can cooperatively inhibit cell proliferation in an AND-logic-gate manner. In addition, drug synergy of PS-ApTCs and ApDCs was evaluated using the Bliss independence model,^68,69^ which showed an average synergistic effect greater than 15%, in a dosage range of 50-400 nM for PS-ApTCs and ApDCs, suggesting a synergistic effect (Figure 4C). A Bliss synergy score was determined as 19.1 (Figure S28), which is comparable to that of previous combination of BET protein degraders and MDR1 inhibitors,^70^ suggesting a synergistic effect. Additionally, an antagonistic interaction was identified between drugs of Control-PS-ApTCs and ApDCs (Figure S29). One potential reason is that Control-PS-ApTCs is inactive in NCL degradation that displays no antiproliferative activity (Figure 2E, 3A, 3B, and S19A). Another is that the competitive internalization process of Control-PS-ApTCs and ApDCs may lead to the inhibition of the antiproliferative effect of ApDCs. Taken together, the above observations confirmed the synergistic antiproliferative effect of PS-ApTCs and ApDCs in HeLa cells. To verify the proliferation inhibition of HeLa cells is really induced by apoptosis, flow cytometry-based apoptosis assay was performed after 48 h incubation of the cells with PS-ApTCs, ApDCs, or PS-ApTCs/ApDCs (Figure 4D). Compared with untreated cells, apoptosis rates induced by above three therapeutic agents increased to 24.1%, 25.8%, and 38.6%, respectively. Meanwhile, early and late apoptotic induction between different treatments was compared (Figure 4E), and, notably, cotreatment of PS-ApTCs and ApDCs induced a much greater degree of late apoptosis than all individual drug-treated groups.

To investigate the biological effect associated with the aforementioned enhanced antiproliferation and pro-apoptotic effects, cell cycle analysis was performed on the same four groups by flow cytometry. As shown in Figure 4F, untreated HeLa cells showed a typical cell cycle distribution pattern with the majority of the cells in G0/G1 phase and a small proportion of approximately 17.4% in G2/M phase. Following treatment with either PS-ApTCs or ApDCs, a dramatic decrease in the G0/G1 phase was observed, while the proportion of cells in the G2/M phase increases to 43.9±3.6% and 55.6±2.7%, respectively. Importantly, PS-ApTCs/ApDCs codosed resulted in a more significant increase in G2/M phase and decrease in G0/G1 phase than individual treatment (Figure 4G), indicating a cooperative strengthening of cell cycle arrest at the G2/M phase by PS-ApTCs and ApDCs.

Based on the remarkable performance *in vitro*, we subsequently investigated the potential mechanisms of their synergistic cytotoxicity. Given that p53 tumor suppressor is a crucial tumor suppressor that promotes cell growth arrest and apoptosis in response to a broad range of cellular damage,^71,72^ we therefore hypothesized that p53 may participate in a critical functional axis of synergistic cytotoxicity. To test this hypothesis, we further performed a quantitative reverse transcription-PCR (qRT-PCR) study to evaluate the p53 mRNA levels in HeLa cells treated with 200 nM of ApDCs, PS-ApTCs, or PS-ApTCs/ApDCs for 48 h (Figure S30). Compared with untreated cells, the mRNA levels of p53 in cells treated with either PS-ApTCs or ApDCs increased similarly by about 1.5-fold (Figure 4H). Notably, cotreatment of PS-ApTCs and ApDCs resulted in a further increase of mRNA level of p53 to about 2.5-fold. These results indicated that the synergistic cytotoxicity of PS-ApTCs and ApDCs might arise from the up-regulation of p53 through two distinct intracellular pathways (Figure 4I). That is, the effective degradation of membrane and cytoplasmic NCL by PS-ApTCs, as confirmed in Figure 2, disables the function of the NCL to p53 mRNA, leading to the up-regulation of p53 in the cytoplasm. On the other hand, the intracellular GSH-activated release of PTX from the ApDCs could impair the function of microtubules, which in turn upregulating the expression of p53. As a result, these two intracellular pathways represent AND-gated logical functions targeting the up-regulation of p53. Taken together, these qRT-PCR data, together with the above *in vitro* cellular results, successfully demonstrate the ability to cooperatively design AND-gated PS-ApTCs and ApDCs for enhanced therapeutic purpose.

### Combinational Delivery of PS-ApTCs and ApDCs Results in Enhanced Synergistic Therapy in Mouse Xenograft Model

The *in vitro* behaviors of the PS-ApTCs and ApDCs motivated us to further investigate their antitumor efficacy *in vivo*. Given that the similar AS1411 aptamer was applied in designing the above two agents, the tumor-targeting properties were first examined using Cy5-modified PS-ApTCs (PS-ApTCs-Cy5) as a model by intravenously administering to HeLa xenografted tumor-bearing nude mice, followed by fluorescence imaging with the IVIS animal imaging system. Free Cy5 and Cy5-modified R-PROTACs (R-PROTACs-Cy5) were applied as the controls. As shown in Figure 5A, the fluorescence signal at the tumor site of the PS-ApTCs-Cy5 treated group increased in a time-dependent manner, and more importantly, with much longer retention time than the control groups. To gain insight into the potential mechanisms of this tumor-targeting property, we further examined the colocalization of PS-ApTCs-Cy5 or R-PROTACs-Cy5 with NCL-expressing cells in the tumor cryosections by immunofluorescence staining using Alexa488-labeled anti-NCL antibody (green), and analyzed with confocal imaging. As shown in Figure S31, more co-localization of Cy5 fluorescent signal with NCL-expressing cells were found in the PS-ApTCs-Cy5 treated mice as compared to the R-PROTACs-Cy5 treated mice. *Ex vivo* imaging was further performed in harvested tumors and major organs at 8 h postinjection (Figure 5B). The fluorescence signal in the tumor tissue collected from the PS-ApTCs-Cy5 treated mice was significantly higher than that from either Cy5 or R-PROTACs-Cy5 treated mice, demonstrating a superior high intratumoral accumulation of PS-ApTCs-Cy5. Moreover, strong fluorescence signals in livers and kidneys while negligible fluorescence signals in hearts and lungs were observed in all treated groups, which were consistent with previous findings.^42^ Taken together, these results demonstrated that the proposed PS-ApTCs and ApDCs possess excellent tumor-targeting capability, which thus promotes the accumulation of PS-ApTCs and ApDCs at tumor sites for subsequent internalization and therapy.

**Figure 5.**
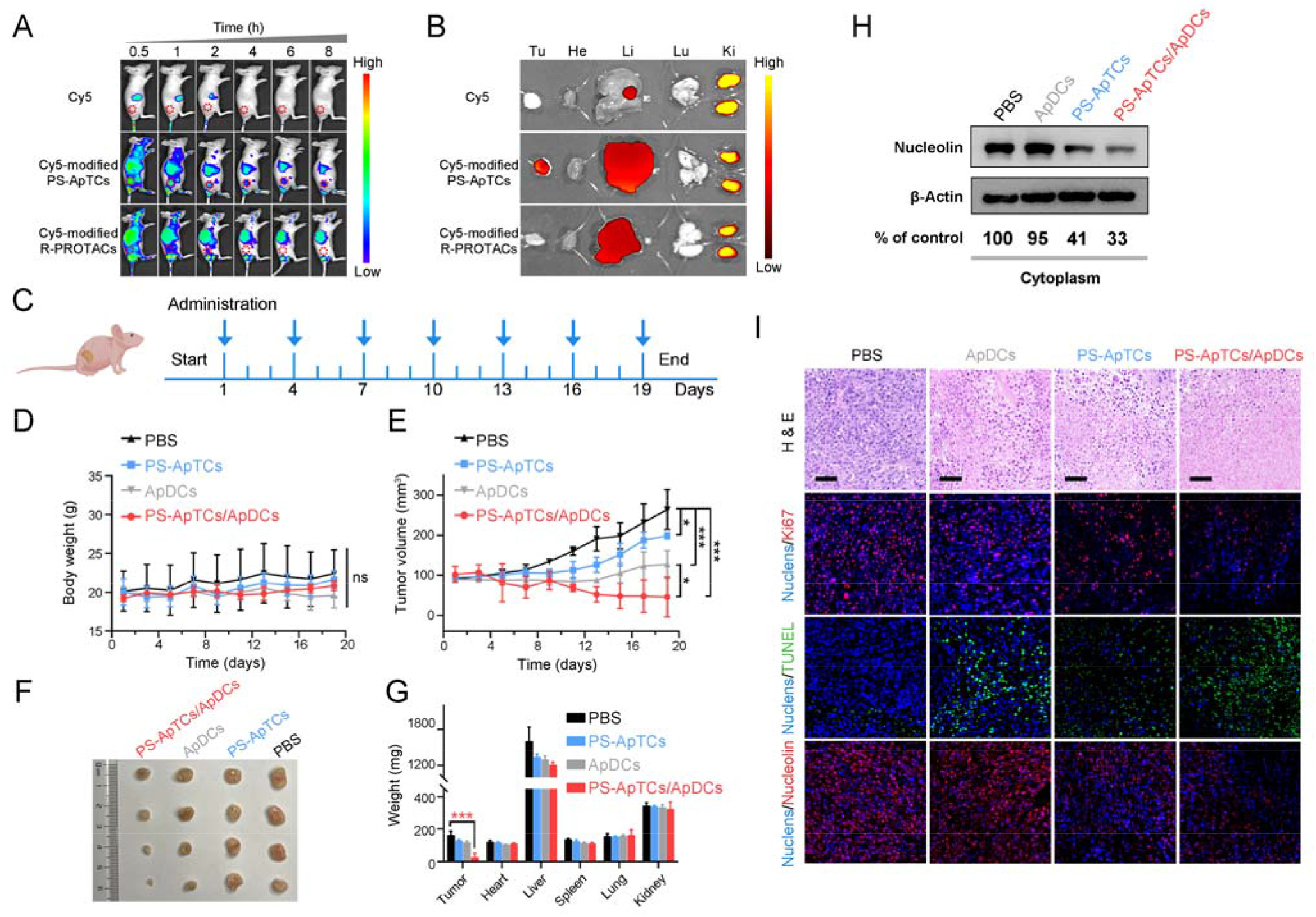
Combinational delivery of PS-ApTCs and ApDCs results in enhanced synergistic therapy *in vivo*. (A) Time-dependent *in vivo* fluorescence imaging showing PS-ApTCs-Cy5 exhibited active tumor-targeting property. (B) Distribution of Cy5, Cy5-modified PS-ApTCs, and R-PROTACs in the tumor (Tu) and major viscera (He, heart; Li, liver; Lu, lung; and Ki, kidney) at organ level 8 h post intravenous injection. (C) Schematic displaying the treatment implemented in the mouse model. The dosing points are indicated with blue arrows. (D) Body weight and (E) tumor volume of mice with different treatment during therapy. (F) Macroscopic views of the xenografted tumor after 19 days of different treatments from the groups indicated. (G) Analysis of tumor and organ weight after 19 days of different treatments at a dosing frequency of every three day *via* intravenous injection. The data are presented as the means ± s.d., n = 4, \**P* < 0.05. ***0.0001 < *P* < 0.001. (H) Expression levels of cytoplasmic NCL in tumor tissue, as determined *via* Western blotting. (I) Microphotographs of H&E, Ki67, TUNEL, and NCL stained tumor tissues from different treatment group. Scale bars, 50 μm.

Next, we evaluated the *in vivo* pharmacokinetic (PK) properties of PS-ApTCs using HeLa xenograft mouse model. After drug administration, the blood concentration of PS-ApTCs was quantified using a ligation-qPCR assay (Figure S32). As shown in Figure S33, PS-ApTCs exhibited a maximum blood concentration (*C*_max_) of 42.82 ± 2.46 nM at 10 min, and was rapidly cleared from the blood and became undetectable within 24 h with a circulation half-life (*t*_1/2_) of 4.44 ± 0.35 h. Other PK parameters (*e*.*g*. AUC, MRT) were also summarized in Table S2. It should be noted that the rapid clearance rate of PS-ApTCs may be attributed to the AS1411-mediated intratumoral accumulation of PS-ApTCs, which has been verified using *in vivo* fluorescence imaging (Figure 5A and 5B). To further evaluate the tumor levels of PS-ApTCs in real time, Cy5-modified PS-ApTCs was intravenously injected to HeLa xenograft mice, and the *in vivo* fluorescence images were collected at indicated time points (Figure S33A). The fluorescence signal at the tumor sites increased gradually in a time-dependent manner (Figure S33B), and the signal intensity peaked at 2 h after drug administration (Figure S33C). Although the tumor levels of PS-ApTCs decreased gradually after 2 h, a stabile fraction remained above 80% in the tumor site after 24 h, which is sufficient to ensure continuous availability of PS-ApTCs over the treatment period. The high tumor accumulation and long tumor retention of PS-ApTCs could be attributed to the tumor-specific targeting ability of AS1411 aptamer (Figure S6) and its good serum stability by PS-modification (Figure 1A). This result is also consistent with the rapid blood clearance rate of PS-ApTCs as mentioned above (Figure S33).

Encouraged by the enhanced tumor accumulation and retention capacity of PS-ApTCs, we further evaluate the *in vivo* therapeutic efficacy of PS-ApTCs and ApDCs using HeLa xenograft mouse model. After the average tumor volume reaches to approximately 100 mm^3^, the mice are randomly divided into four groups (four mice per group) and injected with PBS, PS-ApTCs, ApDCs, and PS-ApTCs/ApDCs, respectively. The treatment process was performed *via* seven tail vein injections over 19 days (Figure 5C), while the body weight and tumor size were monitored every two days. No significant change in the body weight of mice was observed for all the groups (Figure 5D), indicating the low systemic toxicity of these treatments. In contrast, compared with PBS treated groups (Figure 5E), the tumor growth in the PS-ApTCs treated groups was slightly inhibited (*p*<0.05), while ApDCs displayed a significant inhibition of tumor growth (*p*<0.001). In consistency, the tumor size of mice treated with ApDCs for 19 days was reduced to 126.9±34.8 mm^3^, compared to that of 264.2±50.0 mm^3^ for the PBS group and 198.9±8.0 mm^3^ for the PS-ApTCs group (Figure 5F). These results indicated that both PS-ApTCs and ApDCs had a limited tumor growth inhibition effect with a tumor growth inhibition (TGI) of 32% and 79%, respectively, consistent with the *in vitro* data presented in Figure 4. Notably, the tumor growth in the PS-ApTCs/ApDCs group was efficiently inhibited with a TGI of 132% (*p*<0.001), leading to a significant decrease of the tumor weight (Figure 5G, *p*<0.001). This indicated that the combination of the two types of therapeutics was more effective at suppressing the tumor growth than either individuals. In addition, no obvious pathological changes were observed in major organs of the mice after treatment by different groups (Figure S34), which further verify the low toxic side effects of PS-ApTCs and ApDCs. Moreover, our Western blotting results showed that PS-ApTCs or PS-ApTCs/ApDCs codosed could effectively degrade the cytoplasmic NCL in tumor tissues (>50%), compared with the two negative control groups (Figure 5H). Next, we analyzed the tumor pathological changes related to NCL degradation using H&E sections, Ki67 immunofluorescence, and TUNEL staining (Figure 5I). Compared with PBS-treated groups, the H&E staining sections of the PS-ApTCs/ApDCs-treated groups showed obvious nuclear shrinkage and fragmentation, and extensive necrotic areas. However, PS-ApTCs- or ApDCs-treated groups displayed reduced areas of necrosis. In addition, Ki67 immunofluorescence and TUNEL staining results showed that combination therapy of PS-ApTCs/ApDCs displayed the weakest cell proliferation signal but the strongest apoptotic cells signal. Meanwhile, immuno-histochemical analysis revealed that PS-ApTCs- and PS-ApTCs/ApDCs-treated groups significantly downregulated the NCL expression in tumor tissues, while ApDCs-treated groups showed negligible effect. Taken together, the results demonstrated that PS-ApTCs-mediated NCL degradation coupled with ApDCs-based targeted chemotherapy elicited potent antitumor response, indicating enhanced synergistic therapy.

## CONCLUSION

In summary, we developed an effective and cell-type-specific delivery strategy for PROTACs to degrade nucleocytoplasmic shuttling proteins. Using NCL as a model NSP, PS-ApTCs was designed by conjugating phosphorothioate-modified AS1411 aptamer to pomalidomide *via* a simple copper-free click reaction. Taking advantage of both membrane-NCL-mediated internalization and specific recruitment of cytoplasmic NCL and E3, the PS-ApTCs offers significant improvements for spatioselective degradation of NCL in both cell membrane and cytoplasm, resulting in enhanced antiproliferation and pro-apoptotic potencies against target cells. Moreover, we demonstrated for the first time that combination of PS-ApTCs-mediated NCL degradation with ApDCs-based targeted chemotherapy could produce an AND-gated synergistic effect on tumor inhibition. This is particularly attractive when monotherapy has limited anti-tumor efficacy and tends to induce drug resistance. Since *in vitro* selection can obtain aptamers selective for many NCPs, the method demonstrated can expand this versatile PROTACs system significantly to many other NCPs and thus provide a new toolbox to broaden the therapeutic applications of PROTACs. While this manuscript was being prepared, we became aware of a preprint by the Tan laboratory demonstrating an elegant approach by generating a NCL-targeting PROTAC ZL216 using AS1411 and VHL.^73^ Compared to our PS-ApTCs, their strategy focused on the evaluation of tumor-selective degradation of NCL *in vitro* and *in vivo* while the *in vivo* anti-tumor activity as well as potential synergistic effect of their degraders has not been explored. Overall, in addition to highlighting new strategies for PROTACs design, our results provided a starting point for the evaluation of PROTACs-based combinational anti-cancer therapy with the ultimate goal of clinical translation.

## ASSOCIATED CONTENT

### Supporting Information

This material is available free of charge *via* the Internet at http://pubs.acs.org. More detailed experimental section, supporting table S1-S2 including sequences of DNA, and Supporting Scheme S1−S2 and Figures S1−S34.

## AUTHOR INFORMATION

### Author Contributions

All authors have given approval to the final version of the manuscript.

### Notes

The authors declare no competing financial interest.

## ACKNOWLEDGMENT

We greatly acknowledge the financial support from the National Natural Science Foundation of China (no. 22004063), Natural Science Foundation of Jiangsu Province (no. 20200303), and the Fundamental Research Funds for the Central Universities (021414380504). Dr. Xing acknowledges the financial support from National Natural Science Foundation of China (No. 21877032), Hunan Province Talented Young Scientists Program (2019RS2021, 2019RS2023), the Open Research Fund Program of the State Key Laboratory of Analytical Chemistry for Life Sciences (SKLACLS2102), Shenzhen Institute of Synthetic Biology Scientific Research Program (DWKF20210005).

